# Estimating Genetic Inheritance in Case-control Studies

**DOI:** 10.1101/847269

**Authors:** Na Li, Jiayan Zhu, Zhengbang Li

## Abstract

Case-control genetic association study is an efficient tool to search for the deleterious genetic variants predispose to human complex diseases, where the additive mode of inheritance is commonly assumed. However, how the genetic variants influence the occurrence of a certain disease is impossible to know beforehand. We show numerically that the existing procedures using the Hardy-Weinberg equilibrium test to choose the genetic model might be inconsistent. We propose a new method to choose the genetic model by adopting co-dominant code for risk allele and logistic regression model. Extensive computer simulation results demonstrate superiorities of the new method. Applications to six single nucleotide polymorphisms(SNPs) for breast cancer and eight SNPs for Type 2 Diabetes further show good performances of proposed method. In order to specify a genetic model for an allele, our method has some merits and is consistent as sample size is large. We propose to apply our method in related fields.

## 1 Background

The case-control design is an efficient tool for collecting the information of covariate being assumed and tested whether they are associated with human complex diseases in epidemiologic studies. Although it is a retrospective design, the logistic regression model by taking the data as enrolled prospectively is still valid to estimate the coefficients for covariate (Prentice & Pyke, 1979) since the maximum likelihood estimator is consistent and has the asymptotic normality property. Among many covariate such as body mass index, age and blood pressure etc, the genetic variant is one of the most important one. Comparing with insertion, deletion and copy number variation etc, the single-nucleotide polymorphism (SNP), which is a genetic variation that occurs at a specific position in the genome, is a more common form. By now, more than ten thousand SNPs have been identified to be associated with hundreds of human diseases.

When doing an association study, a genetic model needs to be assumed in advance, which refers to a genetic mode of inheritance. Specifying a genetic model means specifying an alternative hypothesis. There are often three commonly used genetic models including recessive, additive and dominant ones. In reality, it is rarely to know the real genetic models. The additive mode of inheritance has been used in many genetic studies (Klein et al., 2005; Hunter et al., 2007; Zheng, Li and Yuan, 2014). However, there are also some other SNPs conferring risk to disease at other modes. For example, Moltke et al. (2014) found a genetic variant p.Arg684Ter associated with 2-h plasma glucose levels and type 2 diabetes at a recessive mode; Nik-Zainal et al. (2016) reported five genes with MED23, FOXP1, MLLT4, XBP1, and ZFP36L1, acting in breast cancer also in a recessive fashion. Mis-specifying the genetic model will result in loss of statistical power. Especially, the genetic model is recessive and the dominant model is adopted and vice visa. So, the Wellcome Trust Case Control Consortium (2007) used the minimum of p-values for score test under the additive model and Pearson Chi-square with 2 degrees of freedom to search for the genetic variant associated with seven common diseases including bipolar disorder, coronary artery disease, crohn disease, hypertension, rheumatoid arthritis, type 1 diabetes, and type 2 diabetes. Sladek et al. (2007) proposed to use MAX (the maximum values of score tests derived under three genetic models) to search for the evidence of SNPs associated with type 2 diabetes. Identifying the genetic model is an challenge problem. As far as we know, there is only work on the basis of Hardy-Weinberg equilibrium test (HWET) to choose the genetic model (Ng and Zheng, 2008; Zheng et al., 2016; Hu et al., 2017). The Hardy-Weinberg equilibrium principle is an important law in population genetics and holds in health individuals of human populations. We will show numerically later that using HWET to discovery the genetic model is inconsistent when the genetic model is additive. This motivates us to develop a new statistical methodology to fill this gap.

In this work, we develop a general framework to infer the genetic model. Our procedure has four merits. Firstly, the methodology is developed in a general setting where a parameter *θ* ∈ [0, 1] is used to represent a genetic model, which not only includes the recessive, additive and dominant models, but also contains other models. It is very suitable for the situation where the imperfect surrogate of the causal SNP is genotyped and the genetic model of the surrogate is derived from the true model of the causal SNP. Secondly, the proposed estimate of genetic model is consistent, which makes up for the shortcoming of using HWET to choose the model. Thirdly, the proposed procedure allows for handling the confounder factors, however, the existing method can not deal with it. Fourthly, we use a binary variable to replace the original genotype values. It can segregate the coefficient of genotype out and make the estimation of parameters feasible.

## 2 Methods

### 2.1 Notations and genetic models

Denote by *Y* the disease status with *Y* = 1 being an individual having a disease and *Y* = 0 being healthy control. Let **X** and *G* be the *m*-dimensional covariate and genotype value, respectively. To detect the relationship between *Y* and *G* with adjusting for the effect of **X**, a typical model is the logistic regression as

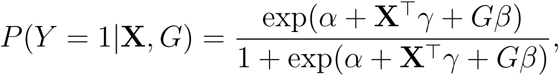

where *α, γ* and *β* are the parameters, and 𝒯 denotes the transpose of a matrix or a vector. We consider a diallelic SNP locus and let the two alleles at a SNP locus be *A* and *a*, and *A* is assumed to be the risk allele or minor allele. Then there are three genotypes as *aa, Aa* and *AA* and the corresponding genotype values are 0, *θ* and 1, respectively, 0 ≤ *θ* ≤ 1. Here *θ* denotes the genetic model. For example, we can set *θ* be 0, 0.5 and 1 for the commonly used recessive, additive and dominant models, respectively.

Suppose that there are *n* subjects including *r* cases and *s* controls randomly drawn from case population and control population, respectively. Let 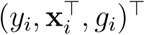 be the observation of the *i*th subject for (*Y*, **X**^⊤^, *G*)^⊤^, *i* = 1, 2, …, *n*. For the sake of simplicity, we assume that the first *r* subjects are cases and the last *s* subjects are controls.

### 2.2 Using HWE Test to choose the model

Denote the genotype frequencies of (*aa, Aa, AA*) in cases and controls by (*p*_0_, *p*_1_, *p*_2_) and (*q*_0_, *q*_1_, *q*_2_), respectively, and let the number of subjects with genotypes (*aa, Aa, AA*) be (*r*_0_, *r*_1_, *r*_2_) in *r* cases and that in *s* controls be (*s*_0_, *s*_1_, *s*_2_). Denote 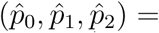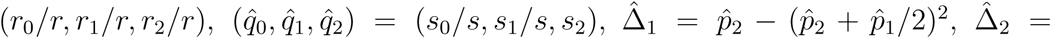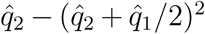, the HWET derived in the whole sample and only in cases can be written as, respectively

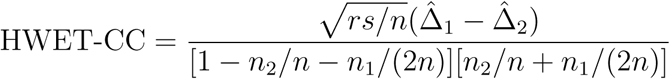

and

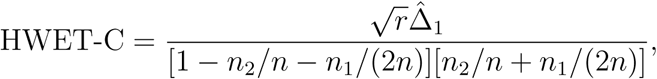

where *n*_1_ = *r*_1_ + *s*_1_, and *n*_2_ = *r*_2_ + *s*_2_. Using the Hardy-Weinberg Equilibrium test to select the genetic model (song et al., 2006; Ng and Zheng, 2008; Zheng et al.,2016) can be summarized as follows: set a positive threshold *c*, for example, *c* = 1.645, the genetic model is determined as: if *Z > c*, the recessive model is selected; if *Z <* −*c*, the dominant model is determined; otherwise, we choose the additive model, *Z* can be HWET-CC or HWET-C. We let the estimate of *θ* be 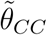 and 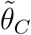 using HWET-CC and HWET-C, repectively. We find numerically that both 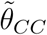 and 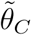 might not be consistent to *θ*.

### 2.3 The proposed procedure

To develop a new method, we decompose the genotype data as

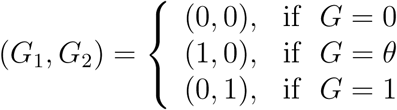

Then, the logistic regression model becomes as follows,

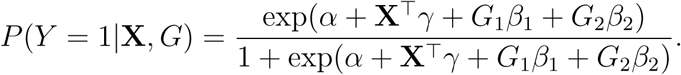

Under rare disease assumption with *α ≪* 0, we can obtain that,

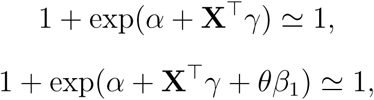

and

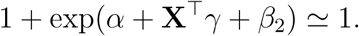

By some algebras, we can obtain that,

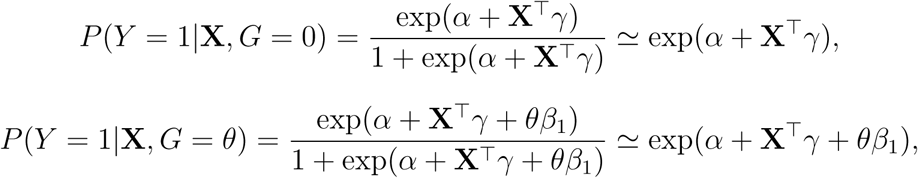

and

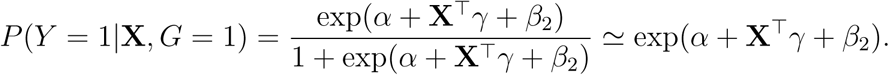

If real genetic model is additive satisfying to

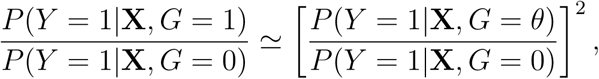

namely,

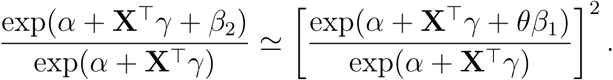

By some algebras, we can obtain that, 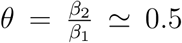. If real genetic model is dominant satisfying to

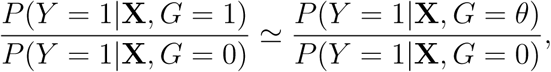

namely,

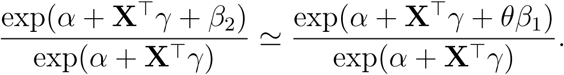

By some algebra, we can obtain that, 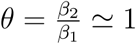. On the basis of above derivation, we can see that, 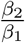 represent genetic inheritance in case-control studies under common recessive, dominant, and additive genetic model. So 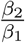 can measure genetic inheritance to some extent in case-control studies.

For *i* = 1, …, *n*, denote observation 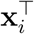 for **X**^⊤^, and (*g*_*i*1_, *g*_*i*2_) for *G*. The likelihood function is

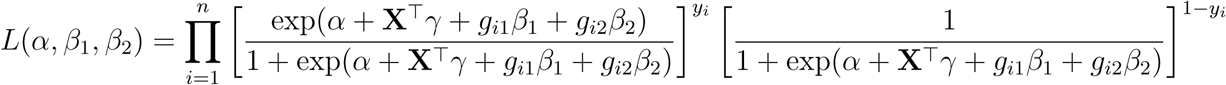

and the log-likelihood function is

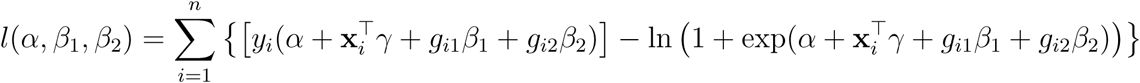

Then the estimate of *β*_1_ and *β*_2_ (denote them by 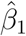 and 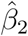, respectively) can be obtained by solving the constrain optimization problem

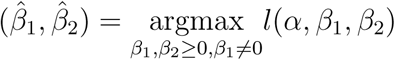

So, the estimate of *θ*, denoted it by 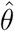, is 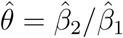. Since 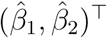 is the consistent estimate of (*β*_1_, *β*_2_)^⊤^ under a general logistic regression setting, so the 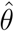 is consistent to *θ*. Or, we can obtain estimate of *θ* directly based on the following log-likelihood function,

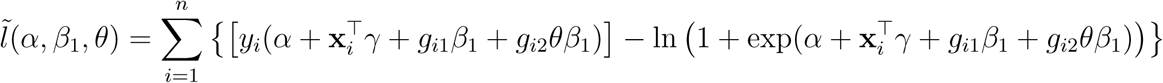

and following constrained optimization problem,

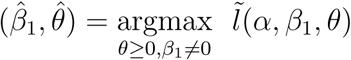

## 3 Results

### 3.1 Simulation Results

To illustrate the performance of the proposed method (denote it by PROPOSED), we conduct simulation studies and compare the mean squared error (MSE) with the existing HWET-CC (Ng and Zheng, 2008) and HWET-C (Zheng et al., 2016) selection procedures. We assume the Hardy-Weinberg equilibrium holds in the general population, that is, the genotype frequencies satisfy to *P* (*aa*) = (1 − *p*)^2^, *P* (*Aa*) = 2*p*(1 − *p*) and *P* (*AA*) = *p*^2^, where *p* = *P* (*A*) and *p* is chosen from {0.05, 0.15, 0.30, 0.45}. Consider two disease prevalences *K* = 0.02 and 0.05, which results in *α* = ln(*K/*(1 − *K*)) = −3.89 and −2.94, respectively, when *γ* = **0**_*m*_ and *β* = 0, where **0**_*m*_ is a *m*-dimensional vector with all the elements being 0. For a fixed *α*, we let *m* = 1 and assume that **X**, independent of *G*, follows the standard normal distribution, *γ* = ln 1.1 and *β* = ln 1.5. *r* = *s* ∈ {500, 1000, 1500, 2000} and *θ* is chosen from {0, 0.25, 0.5, 0.75, 1}. 1000 replicates are conducted to calculate the empirical mean squared error (MSE).

Figures 1 and 2 shows the empirical MSEs of the HWET-CC, HWET-C and PROPOSED for *K* = 0.05. It can be shown that the MSEs of the PROPOSED are decreasing with the sample size increasing. For example, when *r* = *s* = 500 and *p* = 0.15 and the genetic model is recessive, the empirical MSEs of the PROPOSED is 0.18 and that is 0.11 for *r* = *s* = 1000. The empirical MSEs of the HWET-CC and HWET-C under the additive model is almost unchanged for different sample sizes. For example, when *r* = *s* = 500 and *p* = 0.30, the empirical MSEs of the HWET-CC and HWET-C are 0.025 and 0.066, respectively, and those are 0.028 and 0.064, respectively, for *r* = *s* = 1500. As expected, when the sample size is large, the MSEs of the PROPOSED is smaller than those of HWET-CC and HWET-C under the dominant model. For example, when the sample size *r* = *s* = 800 and *p* = 0.45, the empirical MSEs of the PROPOSED, HWET-CC and HWET-C are 0.032, 0.094 and 0.048, respectively.

**Figure 1:**
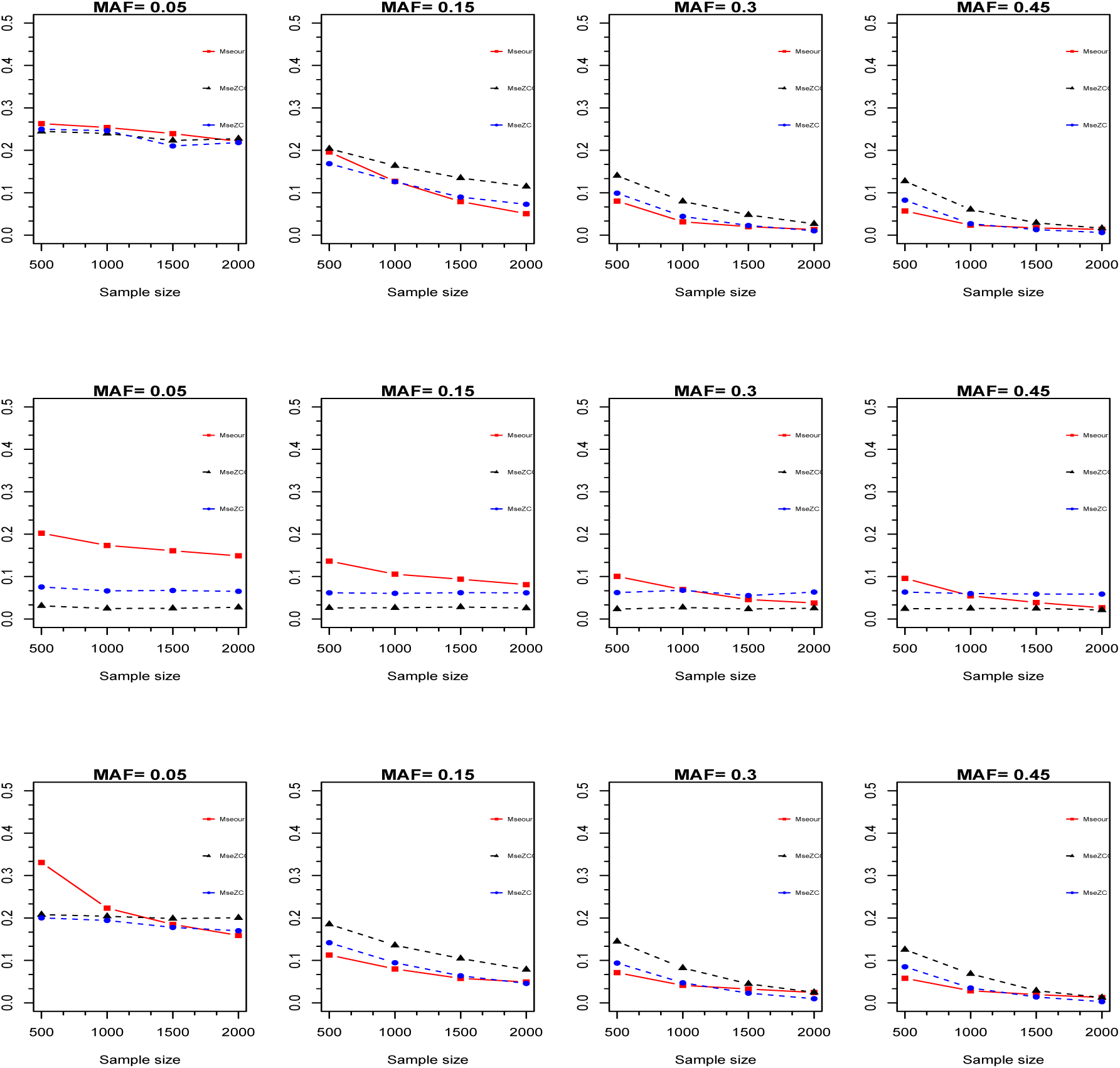
The MSEs of the PROPOSED (square), HWET-CC (triangle) and HWET-C (dot) when *k* = 0.05, *p* ∈ {0.05, 0.15, 0.30, 0.45} and *θ* ∈ {0, 0.5, 1}, where the first row is for *θ* = 0, the second row is for *θ* = 0.5 and the third row is for *θ* = 1. The horizontal axis is the sample size, and the vertical axis is the value of MSEs.

**Figure 2:**
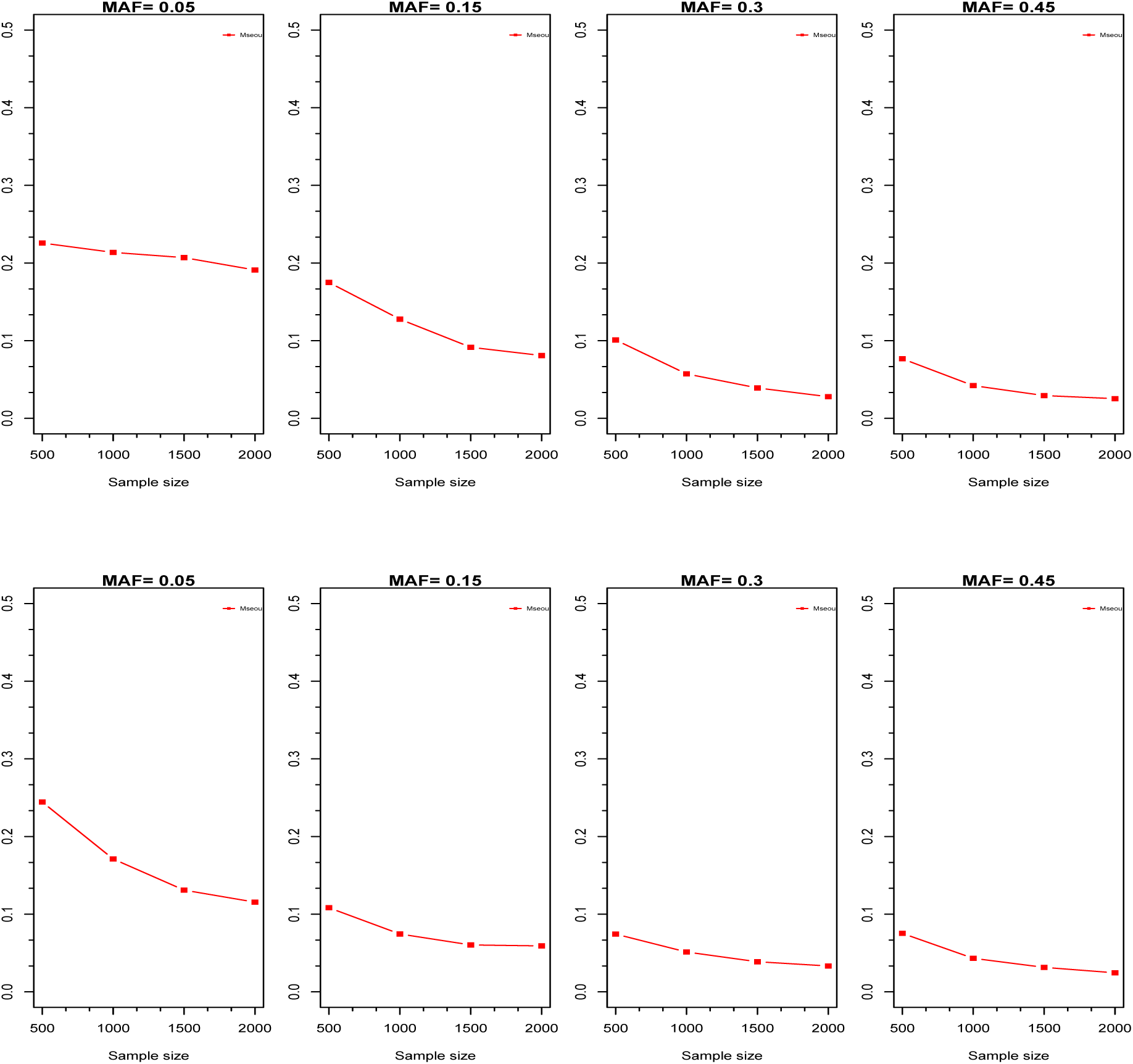
The MSEs of the PROPOSED when *k* = 0.05, *p* ∈ {0.05, 0.15, 0.30, 0.45} and *θ* ∈ {0.25, 0.75}, where the first row is for *θ* = 0.25 and the second row is for *θ* = 0.75. The horizontal axis is the sample size, and the vertical axis is the value of MSEs.

Figures 3 and 4 shows the empirical MSEs of the HWET-CC, HWET-C and PROPOSED for *K* = 0.02. We have similar finds with *K* = 0.05. For instance, when the genetic model is dominant and *p* = 0.30, when *r* = *s* = 1000 and *r* = *s* = 1500, the empirical MSEs of the PROPOSED is 0.081 and 0.040, respectively, and it can be shown that the MSEs of the PROPOSED are decreasing with the sample size increasing. The empirical MSEs of the HWET-CC and HWET-C under the additive model is almost unchanged for different sample sizes. For instance, when *r* = *s* = 500 and *p* = 0.15, the empirical MSEs of the HWET-CC and HWETC are 0.023 and 0.060, respectively, and those are 0.022 and 0.059, respectively, for *r* = *s* = 2000. Again, as expected, the MSEs of the PROPOSED is smaller than those of HWET-CC and HWET-C when the sample size is large under the recessive model. For example, when the sample size *r* = *s* = 1000 and *p* = 0.45, the empirical MSEs of the PROPOSED, HWET-CC and HWET-C are 0.027, 0.065 and 0.035, respectively.

**Figure 3:**
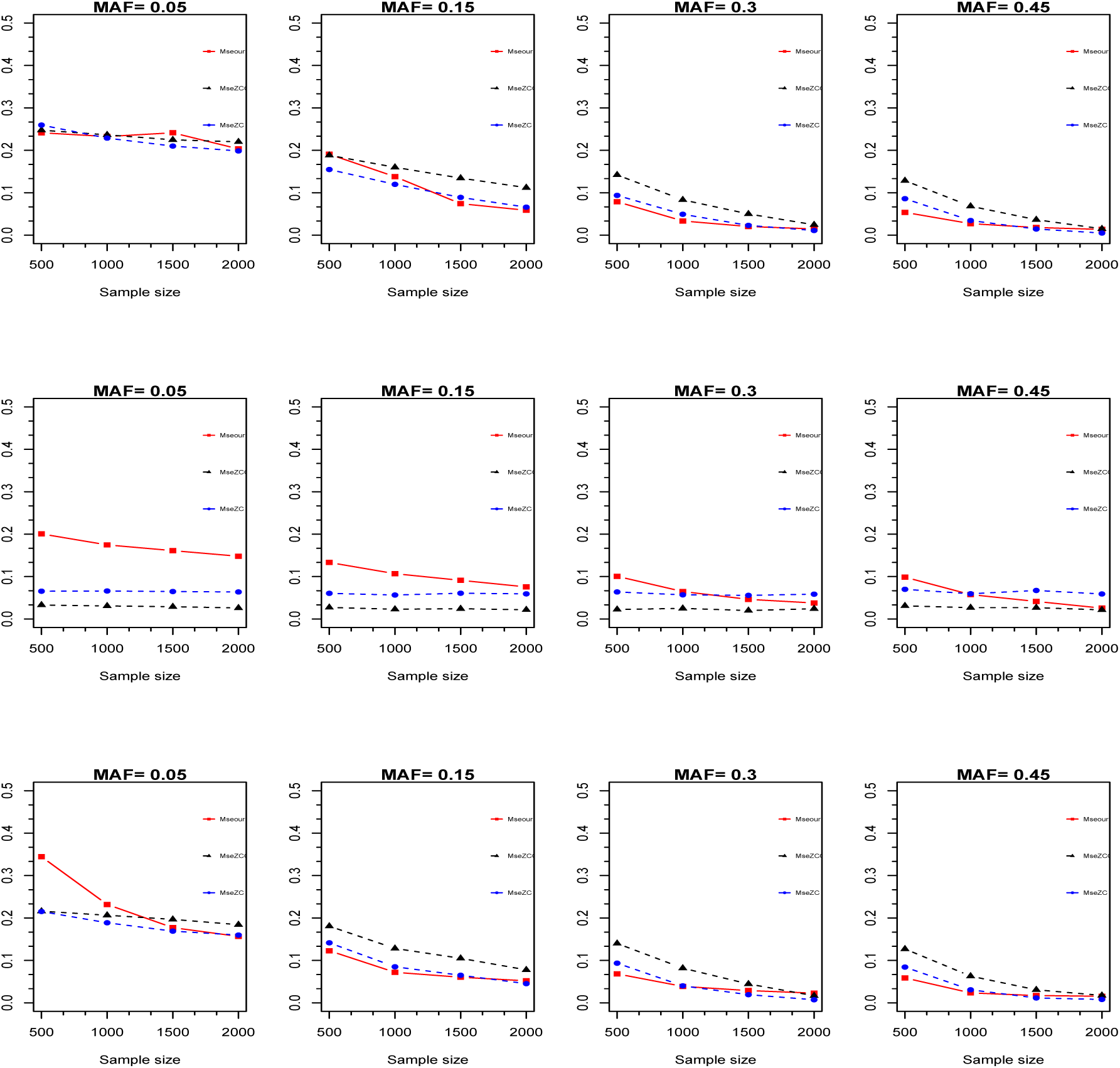
The MSEs of the PROPOSED (square), HWET-CC (triangle) and HWET-C (dot) when *k* = 0.02, *p* ∈ {0.05, 0.15, 0.30, 0.45} and *θ* ∈ {0, 0.5, 1}, where the first row is for *θ* = 0, the second row is for *θ* = 0.5 and the third row is for *θ* = 1. The horizontal axis is the sample size, and the vertical axis is the value of MSEs.

**Figure 4:**
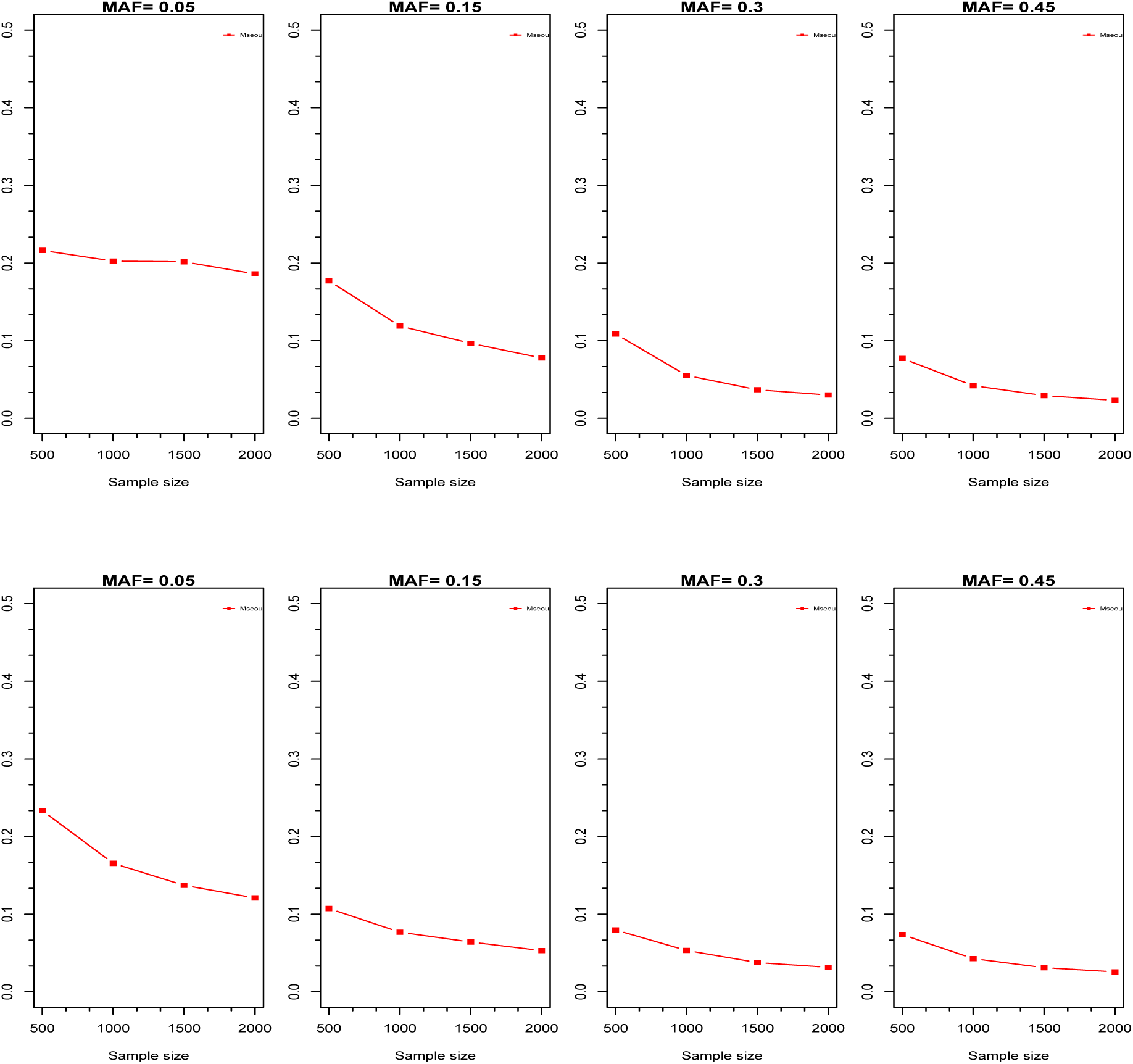
The MSEs of the PROPOSED when *k* = 0.02, *p* ∈ {0.05, 0.15, 0.30, 0.45} and *θ* ∈ {0.25, 0.75}, where the first row is for *θ* = 0.25 and the second row is for *θ* = 0.75. The horizontal axis is the sample size, and the vertical axis is the value of MSEs.

### 3.2 Two Real Application Results

Breast cancer is the common cancer for women. Almost 15 percent of women with breast cancer have family members diagnosed with it, which means that the genetic variants might confer some risk of developing breast cancer. Hunter et al. (2007) conducted a genome-wide association study and have identified 6 SNPs including rs10510126, rs12505080, rs17157903, rs1219648, rs7696175, and rs2420946, associated with breast cancer. The genotype value summaries are shown in Table 1. Type 2 diabetes is a lifelong disease. Typically, the genetic factor confer risk to the type 2 diabetes. Sladek et al. (2007) conducted a genome-wide association study and identified 8 SNPS associated with Type 2 diabetes. The summarized data are also displayed in Table 1.

**Table 1.** The estimated genetic models for 6 SNPs associated with breast cancer and 8 SNPs associated with Type 2 diabetes.

| | $r_0$ | $r_1$ | $r_2$ | $s_0$ | $s_1$ | $s_2$ | $\hat{\theta}$ | $\tilde{\theta}_C$ | $\tilde{\theta}_{CC}$ |
| --- | --- | --- | --- | --- | --- | --- | --- | --- | --- |
| 6 SNPs associated with breast cancer |  |  |  |  |  |  |  |  |  |
| rs10510126 | 955 | 180 | 10 | 854 | 272 | 14 | $2.36 \times 10^{-11}$ | 0.5 | 0.5 |
| rs12505080 | 608 | 477 | 50 | 628 | 408 | 99 | 0.99 | 1 | 1 |
| rs17157903 | 777 | 316 | 18 | 862 | 220 | 26 | 0.99 | 1 | 1 |
| rs1219648 | 352 | 543 | 250 | 433 | 538 | 170 | 0.36 | 0.5 | 0.5 |
| rs7696175 | 353 | 605 | 187 | 396 | 496 | 249 | 0.99 | 1 | 1 |
| rs2420946 | 357 | 546 | 242 | 440 | 537 | 165 | 0.38 | 0.5 | 0.5 |
| 8 SNPs associated with Type 2 diabetes. |  |  |  |  |  |  |  |  |  |
| rs7903146 | 197 | 348 | 149 | 335 | 254 | 65 | 0.62 | 0.5 | 0.5 |
| rs13266634 | 54 | 229 | 411 | 53 | 293 | 307 | $1.62 \times 10^{-8}$ | 0 | 0 |
| rs1111875 | 77 | 302 | 315 | 119 | 308 | 227 | 0.54 | 0.5 | 0.5 |
| rs7923837 | 66 | 300 | 328 | 116 | 296 | 242 | 0.66 | 0.5 | 0.5 |
| rs7480010 | 301 | 327 | 66 | 363 | 246 | 353 | 0.82 | 1 | 0.5 |
| rs3740878 | 25 | 273 | 386 | 65 | 249 | 353 | 1 | 1 | 1 |
| rs11037909 | 25 | 274 | 387 | 65 | 251 | 353 | 0.99 | 1 | 1 |
| rs1113132 | 25 | 271 | 390 | 63 | 251 | 355 | 0.98 | 1 | 1 |

We apply the HWET-CC and HWET-C and PROPOSED to these 14 SNPs to search for their genetic models. The results are given in Table 1. We find that, for the breast cancer, half of them are dominant model and the others are dominant models if the HWET-C or HWET-CC is used. Using the PROPOSED can give the detailed value of the genetic model. For example, for SNP rs10510126, the estimated genetic model is 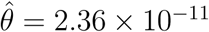, which is recessive model, while using HWET-C gives the additive model. Otherwise, the PROPOSED can give some other model beyond three commonly employed three genetic models. For instance, for SNP rs2420946, using the PROPOSED give a genetic model of 0.38.

## 4 Conclusion

Case-control genetic association study has been proved to be an efficient tool to identify the deleterious variants by scanning the human genome. There are several genetic variants including insertion, deletion, copy number variation, and SNP etc. Among them, the SNP is the most common one. There are 2.96 billion base pairs in human genome and the number of SNPs is about 30 million. By now, more than ten thousand of SNPs have been identified to be associated with hundreds of diseases or traits. To evaluate the significance of a SNP, one has to specify a genetic model. Among three genetic models including recessive, additive and dominant model, the additive model is more frequently to be assumed when conducting an association study. However, in practice, the real genetic model is impossible to know. Especially the causal SNP cannot be genotyped and the SNP locus is its surrogate. Thus the genetic model between the causal SNP and surrogate might be different (Hormozdiari et al., 2015). Misspecifying a genetic model might result in a loss of statistical power.

In this work, we use *θ* to denote the genetic model and *θ* varies range from 0 to 1. The existing work only make the inference for *θ* = 0, 0.5 and 1. It cannot estimate other values of *θ*. By decomposing the genotype score, we proposed a new procedure to estimate *θ*, which is shown to have consistency, while the existing procedures are not consistent based on the numerical results. On the other hand, we obtained the consistent estimate of the genetic model, the next step should be construct the association test based on the chosen model. There is existing the correlation between choosing the genetic model and the association test, which is the future topic.

## Acknowledgement

Prof. Q. Li agrees to withdraw his credit of authorship from the pre-print in the link “https://www.biorxiv.org/content/10.1101/847269v2”.

## Authors Contribution

Jiayan Zhu and Na Li designed the method and conducted simulations, Zhengbang Li wrote the paper and explained all numeric results.

## Data Availability Statement

The raw data supporting the conclusions of this manuscript will be made available by the authors, without undue reservation, to any qualified researcher.

## Compliance with ethical standards

### Funding

This study of Jiayan Zhu was funded by seeding project funding(No. 2019ZZX026), scientific research project funding of talent recruitment, and s-tart up funding for scientific research of Hubei University of Chinese Medicine. This study of Zhengbang Li was funded by the self-determined research funds of Central China Normal University(CCNU) from the colleges basic research of MOE(No.CCNU18QN031).

## Ethical approval

This article does not contain any studies with human participants or animals performed by any of the authors.

